# Exploring cognitive, behavioural and autism trait network topology in very preterm and term-born children

**DOI:** 10.1101/2022.12.09.519321

**Authors:** Marguerite Leoni, Lucy D. Vanes, Laila Hadaya, Dana Kanel, Paola Dazzan, Emily Simonoff, Serena Counsell, Francesca Happé, A. David Edwards, Chiara Nosarti

## Abstract

Compared to full-term (FT) born peers, children who were born very preterm (VPT; <32 weeks’ gestation) are likely to display more cognitive and behavioural difficulties, including inattention, anxiety and socio-communication problems. In the published literature, such difficulties tend to be studied independently, thus failing to account for how different aspects of child development interact. The current study aimed to investigate children’s cognitive and behavioural outcomes as interconnected, dynamically related facets of development that influence one another. Participants were 93 VPT and 55 FT children (median age 8.79 years). IQ was evaluated with the Wechsler Intelligence Scale for Children – 4^th^ edition (WISC-IV), autism spectrum condition (ASC) traits with the Social Responsiveness Scale – 2^nd^ edition (SRS-2), behavioural and emotional problems with the Strengths and Difficulties Questionnaire (SDQ), temperament with the Temperament in Middle Childhood Questionnaire (TMCQ) and executive function with the Behaviour Rating Inventory of Executive Functioning (BRIEF-2). Outcome measures were studied in VPT and FT children using Network Analysis, a method that graphically represents partial correlations between variables and yields information on each variable’s propensity to form a *bridge* between other variables. Results showed that VPT and FT children exhibited marked topological differences. *Bridges* (i.e., the variables most connected to others) in the VPT group network were: SDQ Conduct Problems scale and BRIEF-2 Organisation of Materials scale. In the FT group network, the most important *bridges* were: the BRIEF-2 Initiate, SDQ Emotional Problems and SDQ Prosocial Behaviours scales. These findings highlight the importance of targeting different aspects of development to support VPT and FT children in person-based interventions.

## 1. Introduction

Compared to their full-term born peers (FT; 38-42 weeks’ gestation), children who were born very preterm (VPT; <32 weeks’ gestation) display greater behavioural difficulties such as inattention, emotional dysregulation, socio-communication problems, anxiety, and internalising behaviours (Arpi & Ferrari, 2013; Brydges et al., 2018; Burnett et al., 2013; Johnson & Marlow, 2011; Rothbart et al., 2007). VPT children also exhibit increased autism spectrum condition (ASC)-like traits (Treyvaud et al., 2013) and are more likely to receive an autism diagnosis compared to their term-born peers (7% vs 1.5%, respectively) (Agrawal et al., 2018).

Studies focusing on ASC in VPT and extremely preterm (EPT; <28 weeks’ gestation) samples have revealed a distinct clinical expression from that observed in FT children, suggesting a preterm-specific aetiological ASC profile (Bröring et al., 2018; Jaekel et al., 2013; Verhaeghe et al., 2016). ASC symptoms in VPT/EPT children appear to be primarily rooted in poor socio-emotional abilities and difficulties with social communication and interaction (Jaekel et al., 2013; Johnson et al., 2018; Johnson & Marlow, 2011), rather than a combination of social communication and interactions problems, and rigid and repetitive behaviours and interests, as observed in FT samples. Moreover, social communication and interactions impairments appear to be more homogenously distributed across VPT compared to FT populations, who display greater behavioural heterogeneity (Chen et al., 2019), thus supporting the idea of a preterm-specific ASC aetiology.

VPT children are not only more likely to receive an ASC diagnosis, but also exhibit broader developmental difficulties. A within-group stratification study conducted by Johnson et al. (2018) explored distinct patterns of cognitive and behavioural development in 2 year-old preterm children, and identified a “non-optimal” subgroup characterised by ASC symptomatology, poorer cognitive and language skills, greater social, emotional and attention difficulties, and heightened risk of developing behavioural problems. However, it is unclear how these difficulties identified in VPT children interact or influence one another, highlighting the need to investigate multiple cognitive and behavioural outcomes as a set of interconnected, dynamically interacting facets of development. Indeed, such a paradigm could allow for the identification of core cognitive-behavioural features to better understand preterm-specific ASC symptomatology (Borsboom & Cramer, 2013; Eadeh et al., 2021), in alignment with a (multi-)dimensional and trait-based approach to ASC (Constantino, 2009; Constantino & Charman, 2016; Volkmar & McPartland, 2015).

Previous studies have applied network analysis to ASC populations, studying the network topology of ASC and other behaviours that tend to be comorbid with ASC (Anderson et al., 2015; Montazeri et al., 2019; Ruzzano et al., 2015). These studies have increased our understanding of the dynamic interactions between ASC features and of pathways to outcome phenotypes. Network analysis is a statistical approach which uses nodes (i.e., variables indexing psychological/behavioural constructs) and edges (i.e., partial correlations between nodes) to map and estimate the interaction between psychological constructs. To our knowledge, no prior study has used network analysis to explore multiple facets of development following VPT birth in mid-childhood. In the present study we use network analysis to investigate ASC traits in the context of other behavioural and cognitive measures, in order to understand whether structural relationships between these outcome phenotypes differ between VPT and FT children. We focus on *bridge symptoms*, or nodes connecting different disorders or constructs (Jones et al. 2021), to understand comorbidity and shared characteristics between psychopathology, behavioural, emotional or temperamental difficulties.

We hypothesised that, in addition to VPT participants displaying greater ASC traits, behavioural, cognitive and temperamental difficulties compared to their FT peers, they would show a topologically different cognitive-behavioural network architecture. Specifically, we hypothesised that, similar to findings pertaining to VPT toddlers (Johnson et al., 2018), ASC traits would be more strongly associated with emotional and temperamental difficulties in VPT compared to FT children.

## 2. Methods

### 2.1 Participants

VPT participants, born before 32 completed weeks of gestation, were recruited as new-borns between 2010 and 2013 as part of the Evaluation of Preterm Imaging study (ePrime; EudraCT 2009-011602-42; Edwards et al., 2018), from hospitals within the North and Southwest London Perinatal Networks and were followed up behaviourally throughout childhood (Hadaya et al., 2022; Kanel et al., 2021). The current study focuses on behavioural assessments conducted at a median age of 8.75 years (n=148). Control participants, born after 37 completed weeks of gestation, were recruited from local schools (median age = 8.83 years; n=55). Participants’ demographic and clinical characteristics are presented in Table 1. Exclusion criteria were major congenital malformation, contraindication to magnetic resonance imaging (data not shown here), caregiver(s) unable to speak English or the participant being under child protection proceedings. Ethical approval for the study was granted by London South East Research Ethics Committee (REC: 19/LO/1940) and London Stanmore Ethics Committee (REC: 18/LO/0048).

**Table 1:** Demographic characteristics of the sample:

|  |  | Very Preterm<br>group (n=93) | Control group<br>(n=55) |
| --- | --- | --- | --- |
| Gestational<br>age at birth<br>(weeks) | *Median* | 29.86 | 40 |
|  | *IQR* | [27.57 - 31.86] | [39 - 40.86] |
| Age at testing | *Median* | 8.75 | 8.83 |
|  | *IQR* | [8.33 - 9.17] | [8.25 - 9.17] |
| Sex | *Male : Female* | 50 : 43 | 23 : 32 |
|  | *%* | 53.7%, 46.3% | 41.8%, 58.2% |
| IMD score<br>deciles (N; %) | 1 (least deprived) | 1 | 0 |
|  | 2 | 3 | 2 |
|  | 3 | 10 | 6 |
|  | 4 | 5 | 9 |
|  | 5 | 10 | 2 |
|  | 6 | 11 | 5 |
|  | 7 | 8 | 5 |
|  | 8 | 7 | 3 |
|  | 9 | 12 | 8 |
|  | 10 (most deprived) | 12 | 11 |

### 2.2 Assessment of outcome measures

#### 2.2.1 Cognitive assessment

Participants were administered the Wechsler Intelligence Scale for Children Fourth Edition (WISC-IV; Wechsler, 2003), to assess general intellectual development. The WISC-IV generates a full-scale IQ (FSIQ) score and five subscales evaluating narrower cognitive functioning: *verbal comprehension, perceptual reasoning, processing speed* and *working memory*. Raw scores were transformed into age-normed scaled scores.

#### 2.2.2 Parent-report questionnaires

The participants’ caregiver(s) were asked to complete the following questionnaires. The Social Responsiveness Scale Second edition (SRS-2, Constantino & Gruber, 2012) assessed ASC symptomatology with 65 items comprising two subscales: *rigid and repetitive behaviours* (RRB), and *social communication and interaction* (SCI), matching the two main symptom domains of the latest diagnostic manual for ASC (DSM-V; APA, 2013). Each item is scored along a 4-point Likert scale and raw scores are transformed into T-scores. A score of 76 or higher on the SRS-2 is strongly associated with a clinical diagnosis of ASC (Constantino & Gruber, 2012).

The Strengths and Difficulties Questionnaire (SDQ, Goodman, 2001), assessed child behaviour using five subscales indexing conduct, peer relation problems, emotional problems, prosocial behaviours and a measure of ADHD focusing on hyperactivity-inattention. Responses are given on a three-point Likert scale ranging from 0: “not true” to 3: “certainly true”.

The Temperament in Middle Childhood Questionnaire (TMCQ) for children aged seven to 10 (Simonds & Rothbart, 2004) was administered to measure child temperament. It comprises 157 items and responses range from 1: “almost always untrue” to 5: “almost always true”, with an additional “does not apply” option. These items yield three subscales: *negative affectivity* (NA), *effortful control* (EC) and *surgency* (SU).

Finally, the Behaviour Rating Inventory of Executive Function Second Edition (BRIEF-2; Gioia et al., 2015), a 63-item questionnaire, measured participants’ executive functioning in everyday settings. The BRIEF-2 has nine sub-scale scores: *inhibition, self-monitoring, shift, emotional contro*l, *initiating behaviours, working memory, planning and organising, task monitoring* and *organisation of materials*.

### 2.3 Data analysis

Statistical analyses were conducted using R and RStudio (version 1.3.1).

#### 2.3.1 Univariate group comparisons

Independent samples t-tests were computed to test for differences between VPT participants and FT controls on each subscale score of test or questionnaire administered. All p-values reported are corrected for multiple comparisons controlling the false discovery rate (FDR).

In addition, we tested whether VPT participants had greater ASC-like difficulties (*RRB* or *SCI*) compared to their FT born peers after controlling for sex, age at assessment, and IQ using separate multiple regression models.

#### 2.3.2 Network analysis

The network analyses were conducted using the Gaussian Graphical Model (Lauritzen, 1996) with the *glasso* function, included in the qgraph R package (Epskamp et al., 2012). Separate networks were estimated for the FT and VPT groups, respectively. All subscale outcome measures of interest (verbal comprehension, perceptual reasoning, working memory and processing speed scores from the WISC-IV; inhibition, self-monitoring, shift, emotional control, initiating behaviour, working memory, planning and organising, task monitoring, organisation of material scores from the BRIEF-2; social awareness, social cognition, social communication, social motivation, rigid and repetitive behaviours as measured by the SRS-2; surgency, effortful control and negative affect scores from the TMCQ; and emotional problems, conduct problems, hyperactivity, peer relationship problems, and prosocial behaviour scores measured by the SDQ) were included and equally weighted.

For both the FT and VPT group networks, the following measures were computed for each node: *bridge closeness*, referring to the average distance between one node and any other node that is part of another construct (i.e., questionnaire or assessment); *bridge betweenness*, referring to the number of times a node lies on the shortest path between two other nodes that are from different constructs; *bridge expected influence (EI)*, referring to the sum of all the edges, either negative or positive, between each node and all other nodes from different constructs (Cramer et al., 2010; Eadeh et al., 2021; Epskamp et al., 2012; Jones et al., 2021).

To test group invariance of the two networks regarding their overall structure, *global strength* (i.e., the sum of all edge weights) and specific *edges*, a network comparison test was computed using the *NetworkComparisonTest* R package (Dalege et al., 2017). Given our focus on ASC-like traits, we conducted comparisons between the SRS-2 subscale measures and all other nodes. Edge comparisons across both networks, between SRS-2 subscale measures and all other measures were Bonferroni corrected.

## 3. Results

### 3.1 Sample characteristics and univariate group comparisons

Demographic characteristics of the sample are given in Table 1. There was no significant difference in sex (*χ*^2^(1,141)=1.52, *p*=0.217) or age (*t*(92)=0.23, 95% confidence interval = [-0.2;0.3]; *p* =.822) between the VPT and FT groups.

Mean differences in outcome scores between VPT participants and FT controls are shown in Table 2. In summary, VPT participants had significantly lower IQ and TMCQ effortful control scores, but displayed higher ASC traits, SDQ emotional and behavioural difficulties, and TMCQ negative affectivity scores, and had significantly greater executive function difficulties compared to controls (i.e., BRIEF-2 scores). The mean group difference with the greatest effect size was on ASC-like traits, measured by the SRS-2. Importantly, after controlling for age at testing, sex, full scale IQ, and IMD rank, group differences remained significant for both SRS-RRB (*beta* = 4.46, *t*(118)=2.57, 95% Confidence Interval (CI) = [1.0;7.9]; *p* =.011) and SRS-SCI (*beta*= 5.26, *t*(118)=2.86, 95% CI = [1.6;8.9], *p*=.004). Furthermore, 12 of the 93 VPT participants (12.9%) and 2 of the 55 controls (3.64%) had an SRS-2 score equal to or greater than 76 (*χ*^2^(41,141)=60.7, *p*=0.024).

**Table 2:** Overall and subscale scores on each measure administered and group comparison.

|  | *VPT (N=93)* |  | *FT controls*<br>*(N=55)* |  | *T* | *p* | Cohen's<br>*d* | 95%CI |
| --- | --- | --- | --- | --- | --- | --- | --- | --- |
|  | *Mean* | *SD* | *Mean* | *SD* |  |  |  |  |
| WISC-IV FSIQ | 103.74 | 14.59 | 111.2 | 12.89 | $t(125)=3.22$ | .007** | 0.53 | [2.9; 12.1] |
| VC | 105.36 | 13.37 | 108.31 | 13.8 | $t(110)=1.27$ | .24 | 0.22 | [-1.66; 7.55] |
| PR | 103 | 17.4 | 111.8 | 15.84 | $t(122)=2.95$ | .01* | 0.49 | [2.72; 13.84] |
| PS | 99.42 | 14.86 | 124.33 | 121.25 | $t(55)=1.52$ | .16 | 0.33 | [-8; 57.82] |
| WM | 98.65 | 15.81 | 104.96 | 12.3 | $t(135)=2.7$ | .015* | 0.43 | [1.69; 10.93] |
| SRS-2 | 59.47 | 14.04 | 49.44 | 8.62 | $t(138)=-5.25$ | <.001*** | -0.82 | [-13.8; -6.3] |
| SCI | 54.82 | 11.39 | 47.61 | 8.08 | $t(136)=-4.38$ | <.001*** | -0.7 | [-10.45; -3.95] |
| RRB | 53.85 | 11.41 | 47.26 | 6.57 | $t(138)=-4.35$ | <.001*** | -0.67 | [-9.59; -3.59] |
| SDQ | 11.47 | 5.34 | 9.05 | 4.44 | $t(130)=-2.93$ | .009** | -0.48 | [-4.0; -0.8] |
| Emo | 2.33 | 2.24 | 1.67 | 1.77 | $t(133)=-1.96$ | .069 | -0.32 | [-1.33; .01] |
| Cond | 1.73 | 1.84 | 1.14 | 1.37 | $t(137)=-2.18$ | .044* | -0.35 | [-1.11; -0.05] |
| Peer Rel | 3.16 | 1.66 | 2.53 | 1.33 | $t(132)=-2.55$ | .02* | -0.42 | [-1.14; -0.14] |
| HyperA | 4.23 | 1.93 | 3.7 | 1.94 | $t(114)=-1.49$ | .145 | -0.27 | [-1.18; 0.13] |
| ProSoc | 8.42 | 1.92 | 8.8 | 1.7 | $t(125)=1.21$ | .237 | 0.2 | [-0.24; 0.98] |
| BRIEF-2 | 105.16 | 25.05 | 92.58 | 20.16 | $t(128)=-3.26$ | .004** | -0.54 | [-20.21; -4.94] |
| Beh | 20.46 | 5.73 | 17.19 | 4.97 | $t(121)=-3.56$ | .003** | -0.6 | [-5.09; -1.45] |
| Emo | 25.94 | 7.97 | 23.15 | 6.22 | $t(129)=-2.31$ | .035* | -0.38 | [-5.18; -0.4] |
| Cog | 58.76 | 14.45 | 52.24 | 12.1 | $t(125)=-2.87$ | .01* | -0.48 | [-11; -2] |
| TMCQ | 2.95 | 0.23 | 2.92 | 0.18 | $t(135)=-0.83$ | .409 | -0.13 | [-0.08; 0.04] |
| NA | 2.4 | 0.65 | 2.17 | 0.55 | $t(128)=-2.34$ | .035* | -0.39 | [-0.44; -0.04] |
| EC | 3.28 | 0.45 | 3.52 | 0.4 | $t(127)=3.45$ | .004** | 0.57 | [0.11; 0.4] |
| SU | 3.17 | 0.47 | 3.08 | 0.45 | $t(119)=-1.23$ | .237 | -0.21 | [-0.25; 0.06] |

**Note.** WISC: Wechsler Intelligence Scale for Children; FSIQ: Full Scale IQ; VC: Verbal Comprehension; PR: Perceptual Reasoning; PS: Processing Speed; WM: Working Memory; SRS-2: Social Responsiveness Scale; SCI: Social Communication and Interaction; RRB: Rigid and Repetitive Behaviours; SDQ: Strengths and Difficulties questionnaire; Emo: Emotional Difficulties; Cond: Conduct Problems; Peer Rel: Peer Relationship Difficulties; HyperA: Hyperactivity; ProSoc: Prosocial Behaviours; Int: Initiating Behaviours; Ext: Externalising Behaviours; BRIEF-2: Behaviour Rating Inventory of Executive Functioning, Second Edition; Beh: Behavioural Difficulties; Emo: Emotional Difficulties; Cog: Cognitive Difficulties; TMCQ: Temperament in Middle Children Questionnaire; NA: Negative Affectivity; EC: Effortful Control; SU: surgency.
\*= $p < .05$ ; \*\*= $p < .01$ ; \*\*\*= $p < .001$

### 3.2 Network analysis

#### 3.2.1 Network estimation

Both the VPT (Fig.3) and the FT control group (Fig.2) networks had 26 nodes. Of the 325 possible edges, the VPT network showed 136 (41.84%) that were non-zero, with an overall edge mean weight of 0.02. The FT control group network had 109 non-zero edges (33.54%), with a similar mean weight of 0.02.

#### 3.2.2 Qualitative relationship between variables

Figure 1 displays the FT controls’ network. Visually, social motivation is strongly positively associated with emotional problems and moderately negatively associated with surgency. SRS-2 subscale scores clustered together. Social communication is strongly negatively associated with prosocial behaviours and strongly positively associated with social cognition, but only weakly positively correlated with social awareness. Prosocial behaviours measured by the SDQ are also negatively associated with initiating behaviours and self-monitoring, both measured by the BRIEF-2.

**Figure 1.**
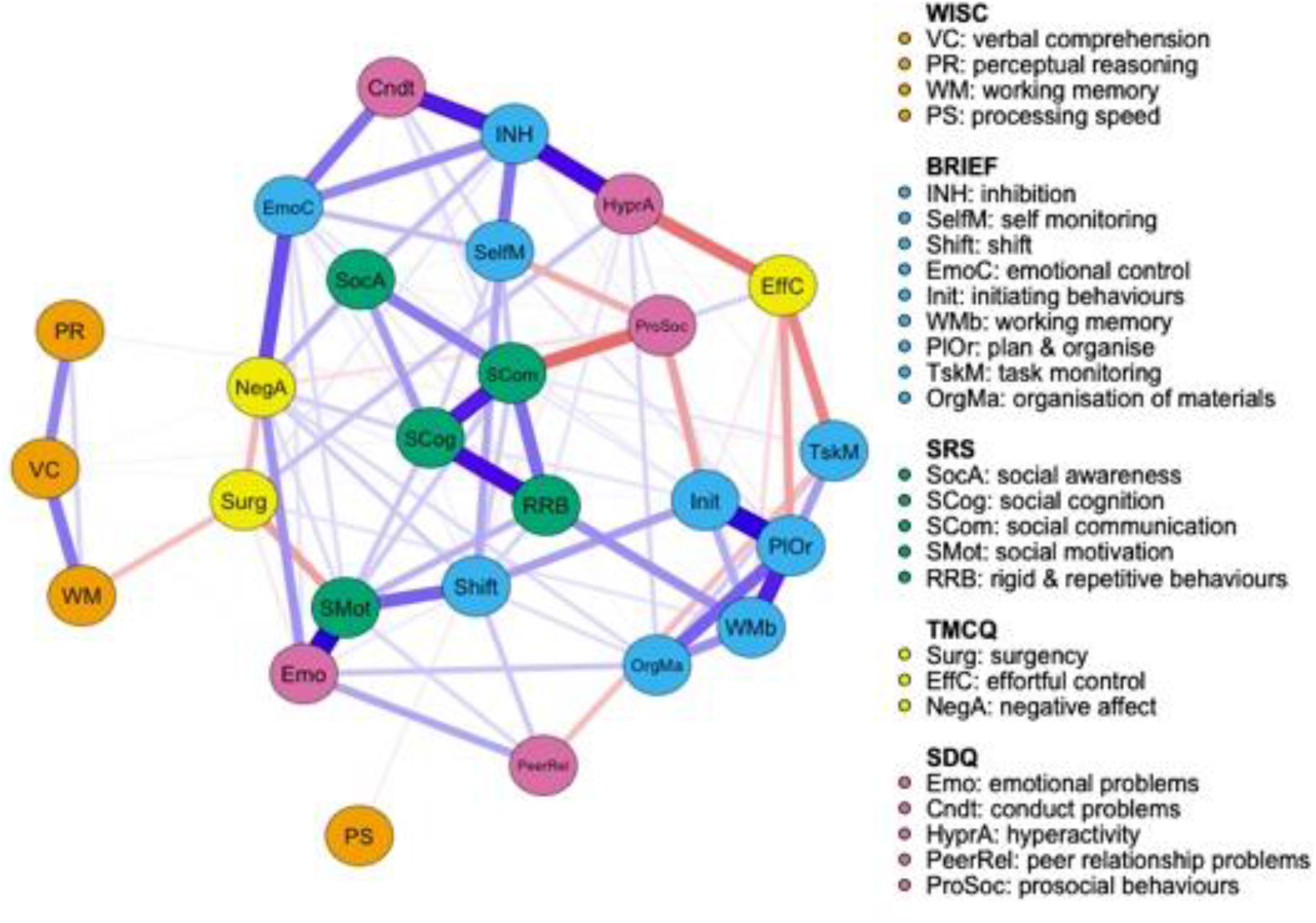
Full term control group network structure of ASC symptomatology and cognitive, executive functioning, temperament and psychopathology outcome variables. ***Note*:** Networks consist of round elements (nodes) which correspond to variables. Lines connecting each node (edges) represent partial correlations between nodes.

Figure 2 presents the VPT participants’ network. Social cognition, as measured by the SRS-2, has a moderate negative association with verbal comprehension, measured by the WISC-IV, and is strongly positively connected to social communication, also measured by the SRS-2. SRS-2 social motivation is strongly negatively associated with surgency, measured by the TMCQ, but positively associated with emotional problems (SDQ), social communication (SRS-2) and attention shift (BRIEF-2). Prosocial behaviours are weak-to-moderately negatively associated with social awareness. SRS-2 subscale scores appear less clustered together in the VPT compared to the FT control group network.

**Figure 2.**
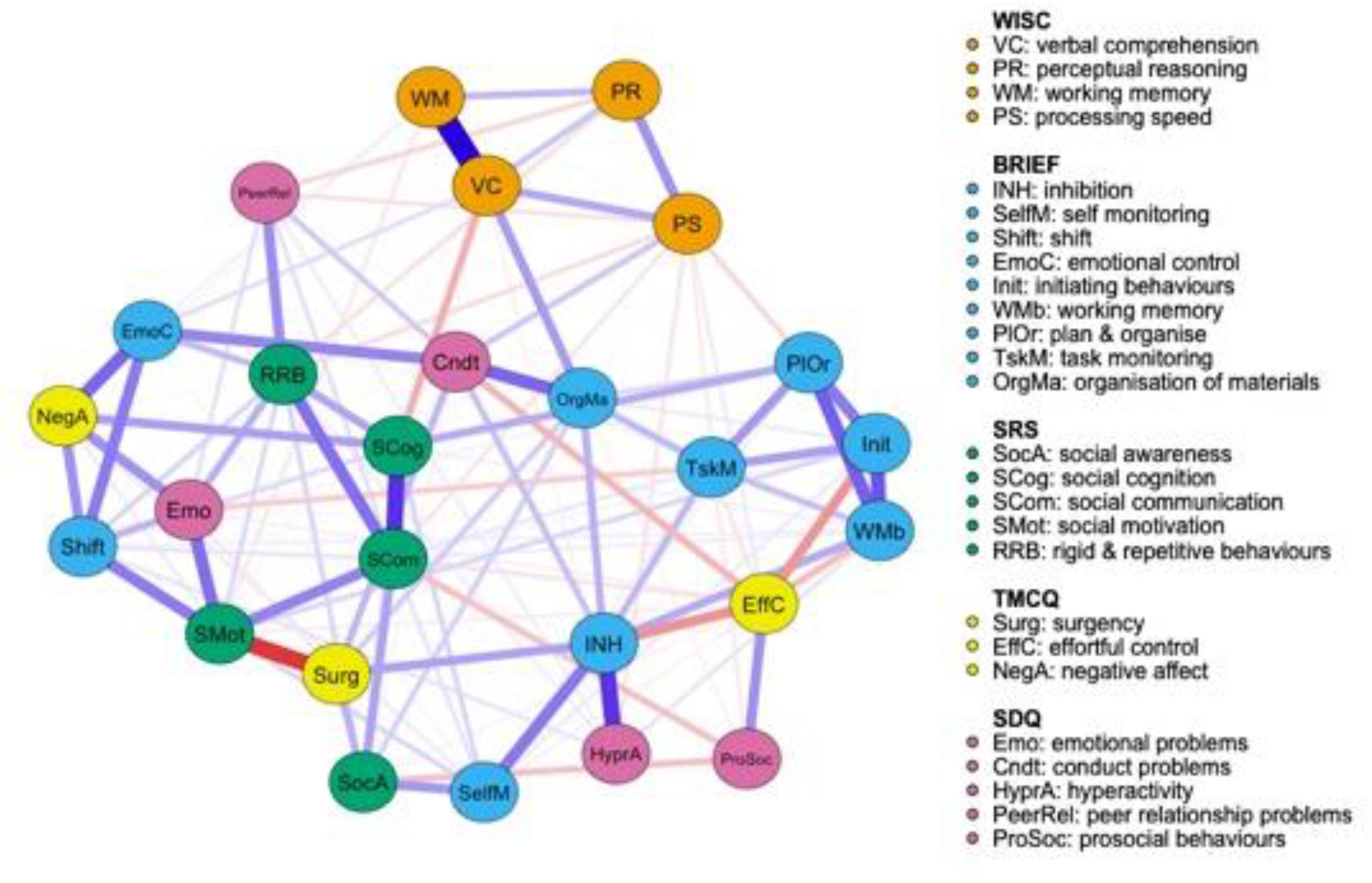
Very preterm group network structure of ASC symptomatology and cognitive, executive functioning, temperament and psychopathology outcome variables.

#### 3.3.3 Node and bridge centrality

Quantitatively, focusing on the bridge centrality of the FT control network, “effortful control” and “prosocial behaviours” had the highest *bridge closeness* scores, “initiating behaviours” and “planning and organising” had the highest *bridge betweenness* values. “Negative affect” and “emotional problems” were the nodes with the highest *bridge expected influence* (Fig.3), suggesting that these nodes were the most important in connecting different behavioural, temperamental, cognitive and ASC-trait constructs.

**Figure 3.**
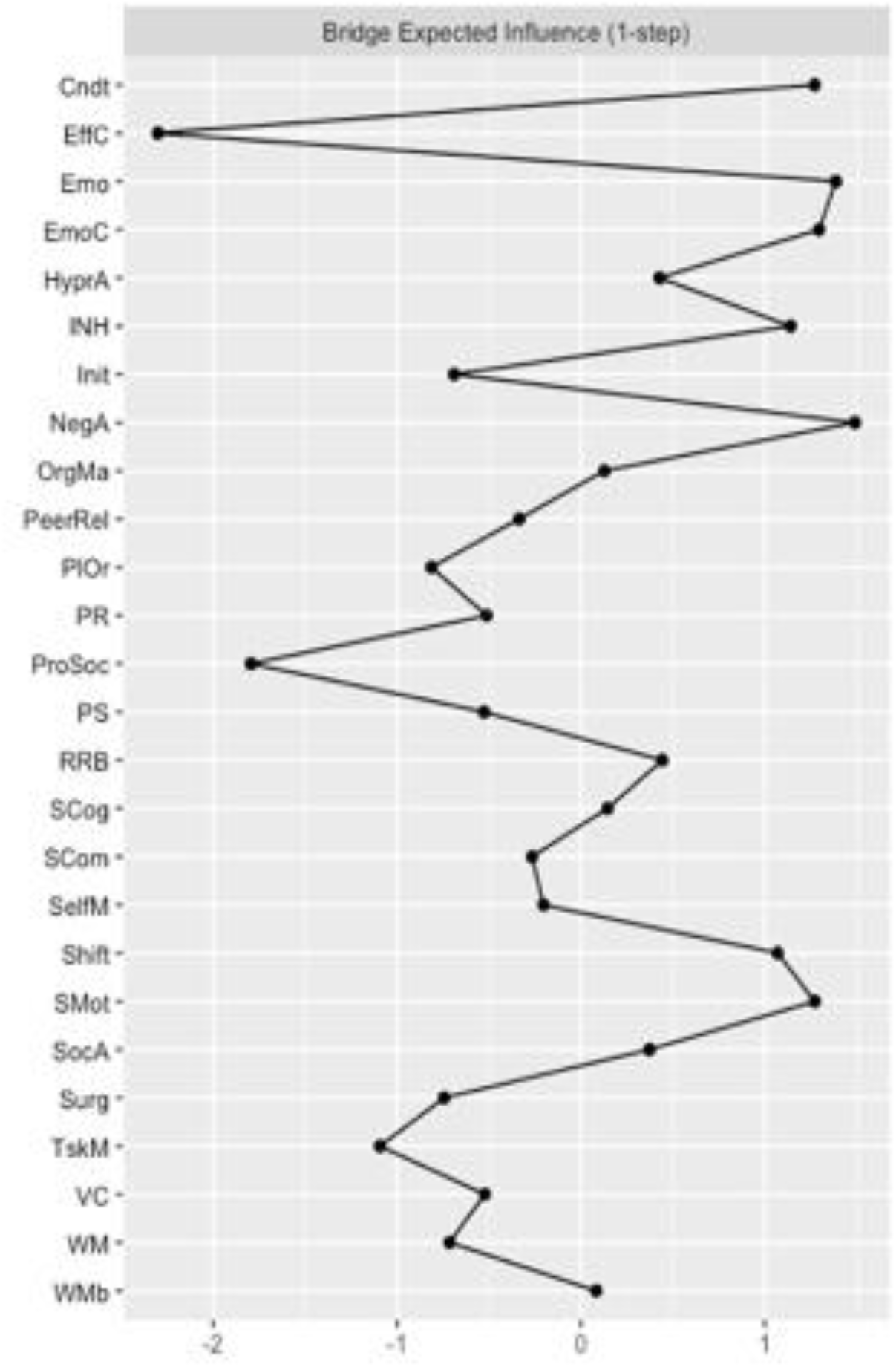
Bridge expected influence for the FT control group network: **Note**. Cndt: conduct problems; EffC: effortful control; Emo: emotional problems; EmoC: emotional control; HyperA: Hyperactivity; INH: inhibition; Init: initiating behaviours; NegA: negative affectivity; OrgMa: organising materials; PeerRel: peer relationship problems; PlOr: planning and organising; PR: perceptual reasoning; ProSoc: prosocial behaivours; PS: processing speed; RRB: rigid and repetitive behaviours; SCog: social cognition; SCom: social communication: SelfM: self-monitoring; SocA: social awareness: Surg: surgency; TskM: task monitoring; VC: verbal comprehension; WM: working memory; WMb: working memory measured by the BRIEF-2.

Regarding the bridge centrality estimates for the VPT network, “organisation of materials” and “conduct problems” were the nodes with the greatest *bridge closeness*. These two nodes were also the ones with the greatest *bridge betweenness* scores. Finally, “negative affect” and “shift” were the nodes with the greatest *bridge expected influence* scores (Fig.4).

**Figure 4.**
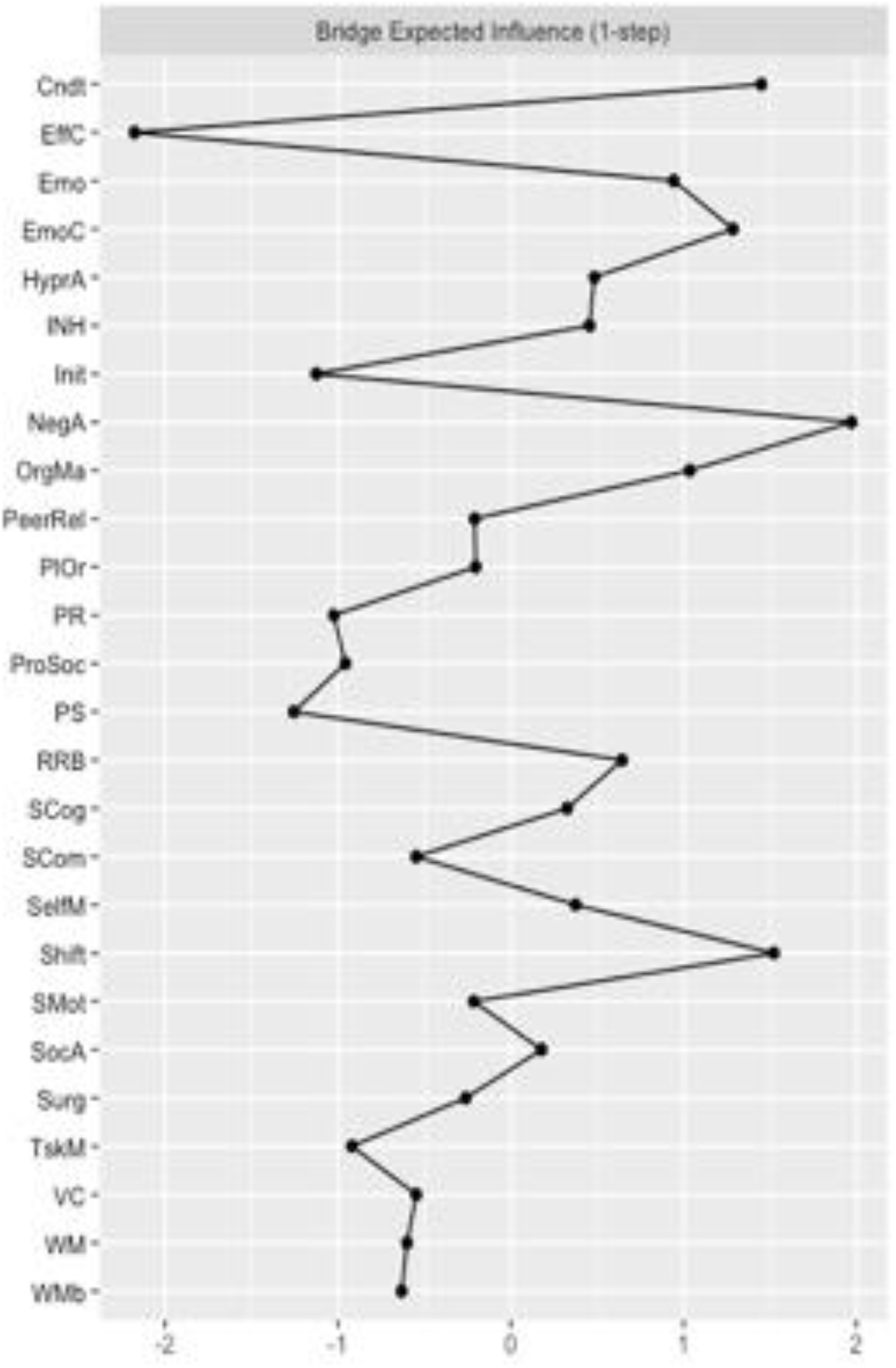
Bridge expected influence for the VPT group network: **Note**. Cndt: conduct problems; EffC: effortful control; Emo: emotional problems; EmoC: emotional control; HyperA: Hyperactivity; INH: inhibition; Init: initiating behaviours; NegA: negative affectivity; OrgMa: organising materials; PeerRel: peer relationship problems; PlOr: planning and organising; PR: perceptual reasoning; ProSoc: prosocial behaivours; PS: processing speed; RRB: rigid and repetitive behaviours; SCog: social cognition; SCom: social communication: SelfM: self-monitoring; SocA: social awareness: Surg: surgency; TskM: task monitoring; VC: verbal comprehension; WM: working memory; WMb: working memory measured by the BRIEF-2.

#### 3.3.4 Network comparison

There was a significant difference in global strength, *p*=.036 and edge weight, *p*=.020 between the overall structure of the VPT and the FT network.

Table 3 presents edge value comparisons (between SRS-2 subscale scores and all other nodes in the networks) that show significant differences between the VTP and FT networks. SRS-2 subscale scores were more connected to other variables in the VPT compared to the FT network. Specifically, VPT participants showed a stronger positive association between “social cognition” and “plan and organise” compared to FT participants. The VPT group also showed a stronger negative association between “social motivation” and “working memory”, as measured by the WISC-IV. The positive connection between “social communication” and “social motivation” was significantly stronger in the VPT compared to the FT network. Finally, the negative correlation between “social motivation” and “surgency” was also significantly stronger in the VPT compared to the FT network.

**Table 3.** Comparison between both networks of significantly different edge values between SRS-2 subscale scores and all other nodes.

| *Edges* | *VPT network* | *FT network* | *p-value for edge* |
| --- | --- | --- | --- |
|  | *Edge value* | *Edge value* | *comparison* |
| Social Cognition – Plan and Organise | 0.14 | 0 | .017* |
| Social Motivation – Working Memory | -0.04 | 0 | .002** |
| Social Communication – Social Motivation | 0.21 | 0.08 | .036* |
| Social Motivation – Surgency | -0.34 | -0.14 | .012* |

*Note.* \* = $p < .05$ ; \*\* = $p < .01$ ; \*\*\* = $p < .001$

## 4. Discussion

In this study we used network analysis to understand how ASC traits, cognitive, behavioural and temperament outcomes dynamically interact in VPT and FT born children. VPT participants displayed elevated ASC traits compared to their FT peers, with group differences in the SRS-2 subscales showing the largest effect sizes of all behavioural measures. Subsequent network analyses allowed us to further investigate the underlying mechanisms that may be driving these behavioural differences. Compared to the FT group, VPT participants’ ASC traits, cognitive, behavioural and temperament outcomes seemed to be predominantly centred around and driven by conduct, executive function difficulties, in particular shifting attention and organising materials, and negative affect. In comparison, the FT participants’ behavioural, cognitive and temperamental profiles appeared to be associated with difficulties in initiating behaviours, emotional regulation, negative affect and to a lesser degree, executive function abilities compared to the VPT children.

The observations in the VPT group reflect the presence of a collection of symptoms commonly reported in VPT populations: poorer socialising abilities, increased internalising behaviours and poorer attention, which are referred to as the “preterm behavioural phenotype” (Johnson & Marlow, 2011). At first glance, the co-occurrence of executive function and socio-communication difficulties seen in the VPT group may be erroneously interpreted as ASC-like social cognition deficits (Alduncin et al., 2014). However, as previously mentioned in the introduction, VPT children may display a unique ASC phenotype, characterised by more predominant difficulties in cognition, attention and socio-emotional processing. Furthermore, despite VPT children exhibiting elevated autistic traits, these do not always reach clinical thresholds (Johnson & Marlow, 2011). Given the isolated and highly clustered SRS-2 subscale scores in the FT network, we tentatively speculate that the SRS-2 in VPT children captures more general behavioural, cognitive, temperamental and socio-emotional difficulties, with a differing etiological pathway to ASC traits or preterm-specific ASC aetiology and symptomatology, rather than more “pure” autistic traits.

In our study, social motivation and social cognition in VPT participants were related to executive functions, in particular working memory, planning and organising. Notably, the VPT network revealed that the link between working memory and social cognition was not direct, but mediated by verbal comprehension. Social cognition deficits seem to impact verbal comprehension, which is in turn associated with working memory. Although the direction of effect cannot be ascertained from our analysis, bidirectional mediating effects might be possible. In a large sample of healthy individuals aged 16 to 91, Froiland & Davison, (2020) found that when controlling for other cognitive measures, working memory had no effect on social perception, or the ability to make inferences and form impressions from social interactions. In their study, verbal comprehension had the largest effect on social perception out of all cognitive measures, with increased verbal comprehension difficulties associated with worse social perception. This finding fits with ASC research in FT adolescents, which highlights poorer verbal comprehension in ASC participants compared to controls, and associated difficulties in social perception (Holdnack et al., 2011). With respect to planning and organising, Vogan et al., (2018) showed that an improvement in planning and organising, self-monitoring and initiating behaviours was associated with reduced social deficits in children with autism. However, improvements in emotional control, shift and inhibition were not associated with better social development. These findings are aligned with our VPT network results, suggesting that VPT children display interactions between ASC traits and other developmental measures that are similar to those observed in ASC clinical groups.

The findings of co-occurring executive function difficulties underlying ASC phenotype and symptomatology in VPT participants is also in line with ASC research in FT, cognitively able children. Pellicano (2010) showed that early executive function skills predicted future abilities in Theory of Mind tasks in children aged four to seven, even when controlling for age and (non)verbal comprehension, thus highlighting the link between socio-cognitive skills and domain-general processes. These findings are also in line with literature revealing associations between cognitive impairments and socio-emotional difficulties in VPT children (Mansson et al., 2014; Rogers et al., 2016). Neuroimaging studies could offer a possible explanation for the overlap between VPT-specific executive function and broader cognitive difficulties and ASC traits, highlighting associations between preterm-related neonatal brain alterations and a later ASC diagnosis (Eklöf et al., 2019; Ure et al., 2016).

Despite no overt between-group difference in surgency scores, reflecting increased shyness and reduced impulsivity, surgency was more strongly associated with less social motivation in VPT compared to FT participants. This result is in line with our previous findings in the same cohort at the age four, which revealed a “social functioning” factor in VPT children reflecting greater emotional, conduct and peer relationship problems and ASC-specific social communication and interaction impairments, as well as poorer prosocial behaviours and lower surgency scores (Kanel et al, 2021). Postulating reciprocal influences between social communication deficits and less social motivation, lower surgency would then reflect more cautious, internalising behaviours.

Our findings highlight the underlying role of executive function and temperament difficulties in the VPT presentation of ASC traits. This is aligned with recent efforts to provide a cognitive understanding of ASC (Frith, 2021; Happé & Frith, 2021), as a unique focus on behavioural phenotypes could lead to a too broad ASC spectrum definition, capturing behaviours that are not intrinsically inherent to ASC. Studies focusing on a cognitive understanding of ASC allow for core differences and specificities to be revealed, which in turn reduces the risk of misattributing VPT phenotype-related behaviours to ASC.

Our findings demonstrate the theoretical and practical relevance of departing from diagnostic-driven labels, and focusing on transdiagnostic traits, such as executive functioning deficits (Krakowski et al., 2020; Siugzdaite et al., 2020; Vaidya et al., 2020). A dimensional approach to understanding psychopathology fits with the concept of equifinality (Cicchetti and Rogosch, 1996), recognising the possible existence of multiple pathways associated with specific outcomes. We have in fact recently shown that only a subgroup of VPT children exhibiting high ASC traits displayed neonatal cerebellar alterations, suggesting distinct aetiological trajectories associated with ASC outcomes (Hadaya et al., 2022). This study, and in particular our results revealing the centrality of executive function and behavioural problems in driving other difficulties for VPT children, further strengthens the argument in favour of person-based treatment approaches targeting underlying behaviours and difficulties rather than surface level symptoms (Kasari et al., 2014). This is particularly important when recognising the potential for these underlying mechanisms, such as poor executive control, to predict later ASC-related difficulties (Pellicano, 2010) and the importance of targeting sub-threshold mechanisms early.

Our study has several limitations. First, since we only considered VPT children, our findings are not generalisable to moderate-to-late preterm children (32-37 weeks’ gestation). Second, the higher prevalence of VPT children reaching a clinical threshold cut-off on the SRS-2, which is parent-rated, compared to other studies, could introduce a rater bias when compared to a clinical observation assessment (Aldridge et al., 2012; Treyvaud et al., 2013). Finally, the weaker associations observed qualitatively between each node, or outcomes, and through the network comparison in the control group, could be due to unequal sample size, and not necessarily linked to core between-group differences.

While this study shows that VPT children are likely at greater risk of displaying ASC traits compared to their FT counterparts, it also highlights that simple group comparisons and focusing on symptom thresholds in isolation, without considering other developmental markers, are not sufficient to fully understand the complexity of the interplay between VPT birth and ASC traits. Our network analysis highlighted underlying constructs leading to ASC behaviours, which may have practical implications, offering a perspective for person-based interventions for VPT children. Indeed, rather than targeting perceived symptoms belonging to specific diagnostic categories, it might be beneficial to focus on their underlying root traits. Future studies can build on these findings by investigating the influence of protective factors that may attenuate the sequelae of VPT birth. We and others have previously shown that a supporting parenting style and a stimulating home environment promoted resilience against behavioural difficulties in VPT samples (Feldman 2007; Ranger et al., 2014; Treyvaud et al., 2009; Vanes et al., 2021; Vinall & Grunau, 2014), and may offer useful avenues of further exploration.

## Acknowledgements

The authors would like to thank the participating families, and research and radiology staff at St Thomas’ Hospital, London involved in this study.

## 5. Declarations

### 5.3 Ethical approval

All procedures performed in this study were conducted in accordance with the ethical standards of the 1964 Helsinki Declaration and its later amendments or comparable ethical standards. The study was approved by Stanmore Ethics Committee (18/LO/0048).

### 5.4 Informed consent

Informed consent was obtained from caregivers and participants.

### 5.5 Author Contribution

Conceptualisation: A.D.E, C.N., M.L. and L.D.V. Methodology: L.H., D.K., M.L. and L.D.V; Formal analysis and investigation: M.L. and L.D.V; Writing – original draft preparation: M.L.; L.H., L.D.V. and C.N.; Writing – review and editing: M.L.; L.D.V.; L.H.; P.D.; E.S.; S.C.; F.H.; A.D.E. and C.N.; Supervision: C.N.; Funding acquisition: A.D.E.; S.C.; P.D.; E.S. and C.N. All authors read and approved the final manuscript.

